# Mosquito Sex Separation using Complementation of Selectable Traits and Engineered Neo-Sex Chromosomes

**DOI:** 10.1101/2025.03.30.646185

**Authors:** Doron SY Zaada, Or Toren, Flavia Krsticevic, Daniella A Haber, Denys Gildman, Noam Galpaz, Irina Häcker, Marc F Schetelig, Eric Marois, Yael Arien, Philippos A Papathanos

## Abstract

Effective and scalable sex separation remains a critical challenge for mosquito genetic control strategies. Genetic sexing strains (GSS) address this by genetically linking maleness with selectable traits, enabling efficient removal of females before release. Here, we describe a robust platform for the development of GSSs in the invasive *Aedes albopictus* mosquito by integrating a CRISPR-engineered selectable phenotype with sex conversion via *nix*, the male-determining factor. As a proof-of-concept, we disrupt the *yellow* gene to generate a vivid pigmentation marker, then rescue its function in males using *nix*-containing transgenes, creating a stable strain where all females are yellow and all engineered males are dark. The resulting GSS males are fertile, robust, and despite lacking the ancestral M locus, exhibit gene expression profiles closely resembling wild-type males. We benchmark sex separation based on pigmentation and discover that *yellow* mutant females exhibit slower larval development, enhancing protandry-based sorting. The GSS strain is compatible with existing size-based sex sorting systems, allowing for improved separation accuracy through the integration of natural and engineered sexually dimorphic traits. Additionally, we find that GSS females lay desiccation-sensitive eggs, reducing the risk of accidental female releases. Our approach is the first to engineer a sex-linked selectable trait by precisely targeting an endogenous gene and restoring its function in males, establishing a versatile platform for GSS development in *Aedes* mosquitoes.

## Introduction

*Aedes albopictus*, the Asian tiger mosquito, has emerged as one of the world’s most invasive disease vectors, capable of transmitting arboviruses such as Dengue, Zika, and Chikungunya (Nie & Feng 2023). Traditional control strategies, including insecticide application and habitat management, have struggled to contain mosquito populations, driving increased interest in genetic control approaches (Alphey 2014; Raban *et al*. 2023). Genetic control strategies have now been developed based on a number of different underlying technologies, including irradiation-based sterilization of the Sterile Insect Technique (SIT), or more recently using symbionts, genetic modification and gene drives (Wang *et al*. 2021). Most mosquito control strategies rely on the mass release of males, necessitating the complete removal of females. Released females can spread diseases, increase nuisance biting and interfere with the dispersal of released males. In *Wolbachia*-based suppression programs, female releases can compromise the sustainability of control programs. However, existing sex-separation methods, such as pupal size sorting, remain inefficient, costly, and difficult to scale for large-scale mosquito control programs (Balatsos et al. 2024; Crawford et al. 2020).

Genetic sexing strains (GSS) provide a scalable and cost-effective solution by linking maleness to a selectable trait, allowing for the efficient or automated removal of females. A GSS typically consists of at least two components: (1) an autosomal recessive mutation that disrupts a visible or conditional selectable trait, and (2) linkage of a wild-type (WT) rescue allele of this gene to the male-specific chromosome, ensuring trait restoration exclusively in males (Augustinos *et al*. 2017; Franz 2005). Historically, the development of a GSS in mosquitoes has been hampered by the laborious and unpredictable nature of classical mutagenesis and translocation approaches. Recent advances in CRISPR/Cas9 gene editing offer a predictable engineering approach, enabling precise, targeted modifications.

*Aedes* mosquitoes lack morphologically distinct sex chromosomes and sex is determined by a small, non-recombining autosomal region known as the M-locus (Newton *et al*. 1974). Within this region the male-determining factor *nix* functions as a master regulator of sex determination and defines male development (Hall *et al*. 2015). Engineered insertions of *nix* can function as synthetic neo-sex chromosomes, converting genetic females into fertile pseudomales and resulting in the loss of the ancestral M-locus in these populations (Lutrat *et al*. 2022; Zhao *et al*. 2022). We reasoned that *nix*-mediated sex conversion could be leveraged to develop a robust GSS based on linking selectable rescues to masculinized pseudomales in a mutant background.

Here, we demonstrate this approach by engineering a GSS in *Ae. albopictus* that integrates CRISPR-disruption of an endogenous gene resulting in a visible phenotype (*yellow*) with *nix*-mediated male rescue. The *yellow* gene, a well-characterized melanization factor, produces a viable and vivid mutant phenotype in *Ae. aegypti* and *Ae. albopictus* (Li *et al*. 2017; Liu *et al*. 2018). We disrupted *yellow* to generate a stable, homozygous mutant strain in which all mosquitoes display the *yellow* phenotype. We then restored *yellow* function with engineered rescues containing both mini-*yellow* and *nix*, yielding a pure-breeding GSS where all males are dark and all females remain yellow.

We evaluate this GSS in terms of its stability of sex-conversion, impact on sex-biased gene expression, and practical utility for sex separation. We find that pseudomales exhibit gene expression profiles closely resembling WT males, confirming successful sex conversion. We also show that GSS females display delayed larval development, enhancing protandry-based sex sorting, and produce desiccation-sensitive eggs, which could contribute to genetic containment of unintentionally released females in field applications. This work provides a versatile platform for engineering GSSs in *Aedes* mosquitoes and highlights a path forward for the development of mosquito population suppression strategies.

## Results

### Generating a *yellow* mutant as a selectable trait

To develop a genetically encoded sex-separation marker, we used CRISPR/Cas9 to generate a *yellow* mutant strain in *Ae. albopictus*. The *yellow* gene is located on chromosome 1, approximately 81.7 Mb from the M-locus. It has a relatively simple two-exon structure, producing a single mRNA isoform (**Fig. 1A**). We targeted the first exon with sgRNA^134^ and injected pre-assembled Cas9-RNP complexes into syncytial embryos. High rates of injected mosquitoes displayed mosaic pigmentation and those surviving to the adult stage were backcrossed to WTs to establish isofamilies. As low recombination rates were expected between the *yellow* locus and the nearby non-recombining region of chromosome 1, we intercrossed F1 adults from different isofamilies to obtain F2 homozygous *yellow* mutants, which were crossed to establish a stable mutant strain. As expected from the intercrossing of F1s, amplicon sequencing revealed four different circulating null alleles, all containing frameshift mutations that resulted in premature stop codons before the *yellow* Major Royal Jelly Protein (MRJP) domain (**Fig. 1B**).

**Figure 1.**
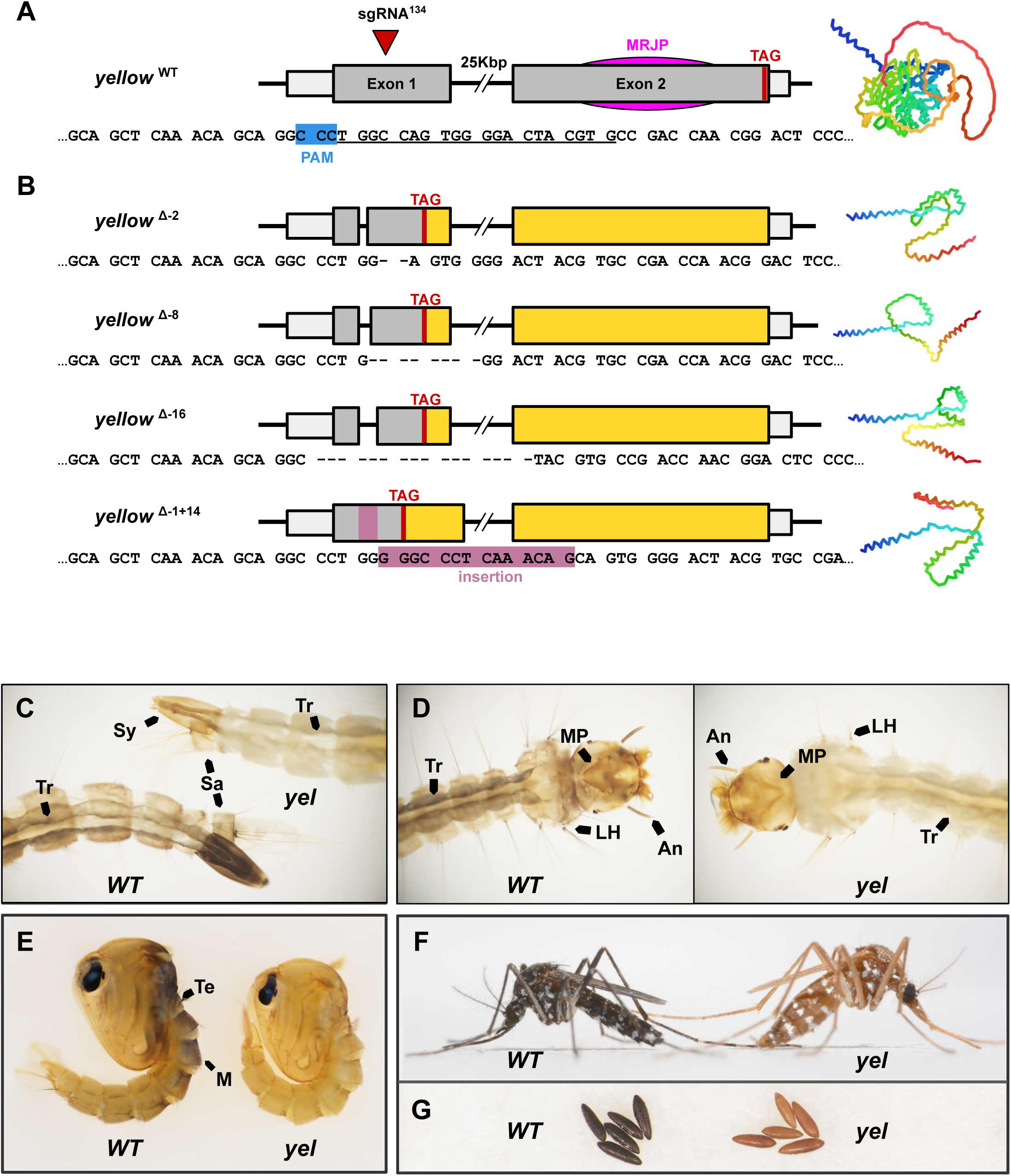
Characterization of the *yellow* mutant phenotype. **A)** Structure of the *yellow* gene showing the target of sgRNA^134^ in the first exon, its target sequence (underlined) and PAM site (blue), the Major Royal Jelly Protein (MJRP) domain and premature TAG stop codons. AlphaFold2-predicted 3D structure of *yellow* is shown on the right. **B)** CRISPR-induced loss-of-function *yellow* mutant alleles isolated. All mutations cause a frameshift leading to premature stop codons and protein structures lacking the MJRP domain. **C-G)** Comparison of *WT* and *yellow* phenotype across development indicating pigmentation differences in key structures with arrows: **C,D)** Fourth instar larvae - trachea (Tr), syphon (Sy), saddle (Sa), lateral hairs (LH), mental plate (MP), and antenna (An). **E)** One-day-old male pupae - tergum (Te) and mesothorax (M). **F)** Adult females. **G)** One-day-old eggs laid by WT and *yellow* females.

Consistent with the role of *yellow* in melanin biosynthesis(True 2003), mutants failed to properly deposit melanin, resulting in a striking pigmentation phenotype throughout development (**Fig. 1C-G**). Larvae displayed reduced cuticular melanin, particularly in antennae, mental plates, lateral hairs, trachea, syphons and saddles. (**Fig. 1C-D**). In pupae, pigmentation differences were most pronounced in the dorsal mesothorax and tergum, irrespective of pupal age (**Fig. 1E**). In adults the characteristic black-and-white “tiger stripes” of *Ae. albopictus* appeared yellow (**Fig. 1F**). Eggs laid by *yellow* females failed to fully melanize, remaining golden-brown instead of black as WT eggs (**Fig. 1G**), consistent with previous studies (Noh *et al*. 2021). Notably, this egg melanization phenotype was maternally inherited and was not restored by crossing *yellow* females to WT males.

### Transgenic rescue of the *yellow* phenotype

To restore pigmentation, we designed a minimal *yellow* (mini-*yellow*) rescue construct that removed the large 25-kbp intron between the two exons. To do so, 1.1-kbp of upstream regulatory regions followed by exon 1 were amplified and cloned in-frame with exon 2 flanked by 660bp of presumed terminator (**Fig. 2A**). This mini-*yellow* rescue, called EnYR, was cloned into a *piggyBac* transformation construct also containing an eGFP transformation marker, driven by OpiE2 (**Fig. 2A**). Given the untested nature of these regulatory regions, we also constructed a second mini-*yellow* rescue, called HspYR, where the mini-*yellow* was driven by the ubiquitous *Ae. albopictus hsp83* promoter, previously validated using a DsRed reporter (**Fig. S1**).

**Figure 2.**
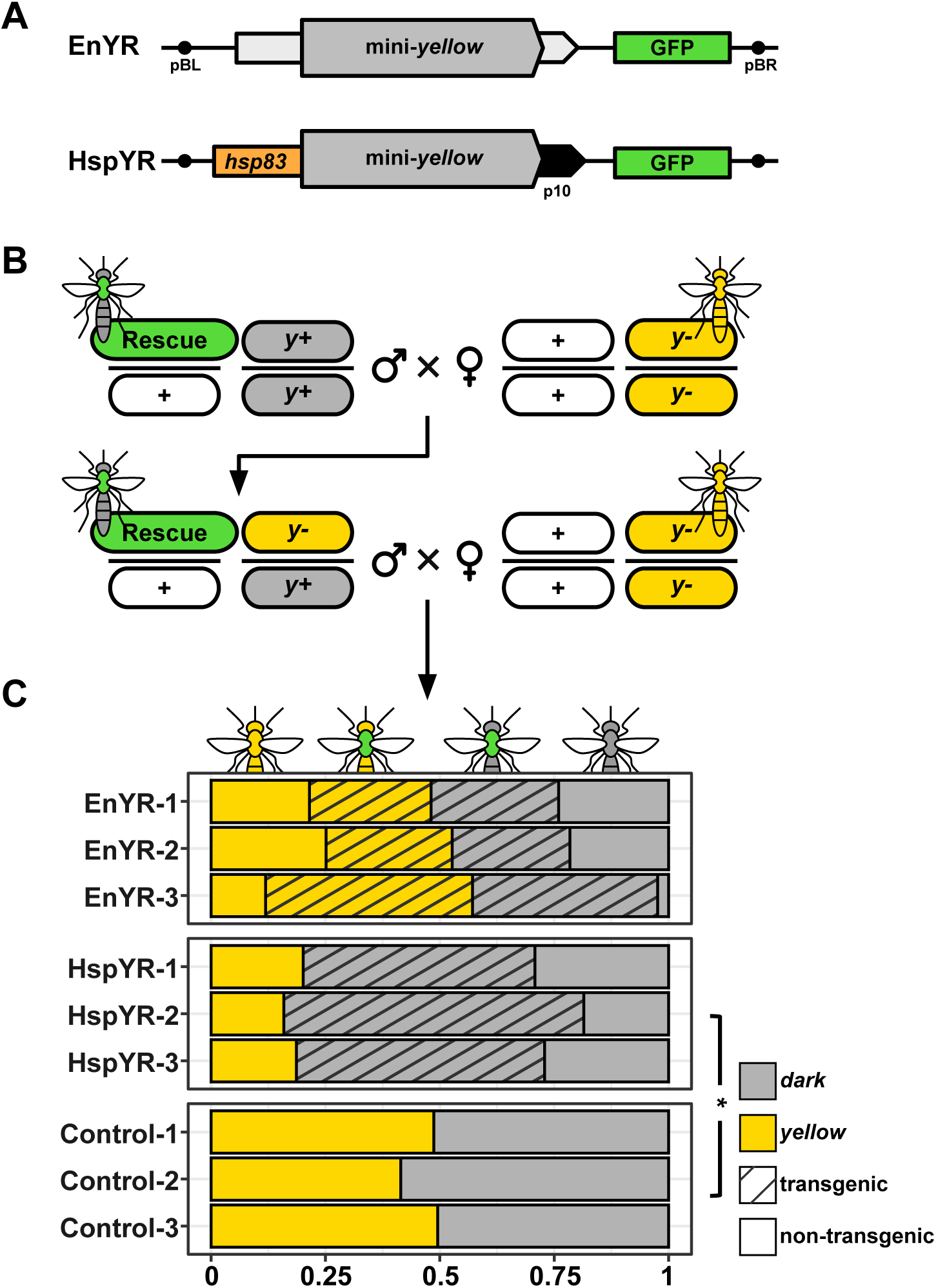
Transgenic mini-*yellow* restores mosquito pigmentation. **A)** Structure of EnYR and HspYR rescue constructs containing the mini-*yellow* coding sequence (CDS) driven either by endogenous *yellow* regulatory elements or the *hsp83* promoter and p10 terminator. Both constructs also contain the OpIE2:eGFP transformation marker and *piggyBac* arms. **B)** Crossing scheme to test complementation. Transgenic males homozygous for the native *yellow* (*y^+^*) allele were crossed to homozygous *yellow* (*y^-^*) mutant females, producing F1 progeny heterozygous at the *yellow* (*y*^+^/*y*^-^) locus and carrying the mini-*yellow* rescue construct. F1 transgenic males were then test-crossed with homozygous *yellow* (*y^-^*) mutant females to evaluate pigmentation restoration in F2 progeny. As a control non-transgenic F1 heterozygous males were crossed to homozygous *y^-^*females. **C)** Distribution of pigmentation phenotypes for dark and *yellow* among transgenic and non-transgenic individuals. Significant reduction from a default 50% *yellow* (Z-test, p < 0.05) is indicated.

Microinjections of eggs of the *yellow* strain with the mini-*yellow* resulted in extremely low post-injection survival (see below). We therefore injected WT eggs and introgressed transgenes into the *yellow* background. Three independent transgenic strains were established for each mini-*yellow* construct and transgenic males from each of the six isofamilies were sequentially crossed to *yellow* females for two generations (**Fig. 2B**).

In F2, we evaluated the overall frequency of the *yellow* phenotype and its co-occurrence with the eGFP marker, which would indicate lack of genetic rescue from the transgene. In crosses involving the EnYR construct, both yellow and dark progeny containing the transgene (eGFP-positive) were observed and the overall frequency of the two phenotypes was similar to control crosses, involving non-transgenic F1 males (**Fig. 2C**). These results suggested that EnYR failed to rescue pigmentation. In contrast, HspYR fully restored pigmentation, with all transgenic individuals displaying WT-like dark coloration (**Fig. 2C**). The proportion of dark F2 individuals increased to 81.78 ± 1.24%, indicating successful genetic rescue in all lines. These results confirmed that the *yellow* regulatory regions we selected failed to correctly express mini-*yellow,* whereas the *hsp83* promoter drove sufficient expression for effective *yellow* complementation.

### Combining sex conversion and *yellow* rescue

To genetically associate mini-*yellow* rescue to maleness, we next inserted *nix* within HspYR, generating construct YRN that was injected into WT embryos **(Fig. 3A)**. The *nix* isoform we used was previously shown to transform genetic females into fertile males, which are referred to as pseudomales from here (Lutrat *et al*. 2022; Zhao *et al*. 2022). Seven independent transgenic isofamilies (YRN1-7) were established from independent founders. Males or pseudomales from each isofamily were individually backcrossed to *yellow* females for multiple generations until a pure-breeding sexing strain was isolated producing exclusively dark, eGFP-positive males and yellow, eGFP-negative females **(Fig. 3B-D)**.

**Figure 3.**
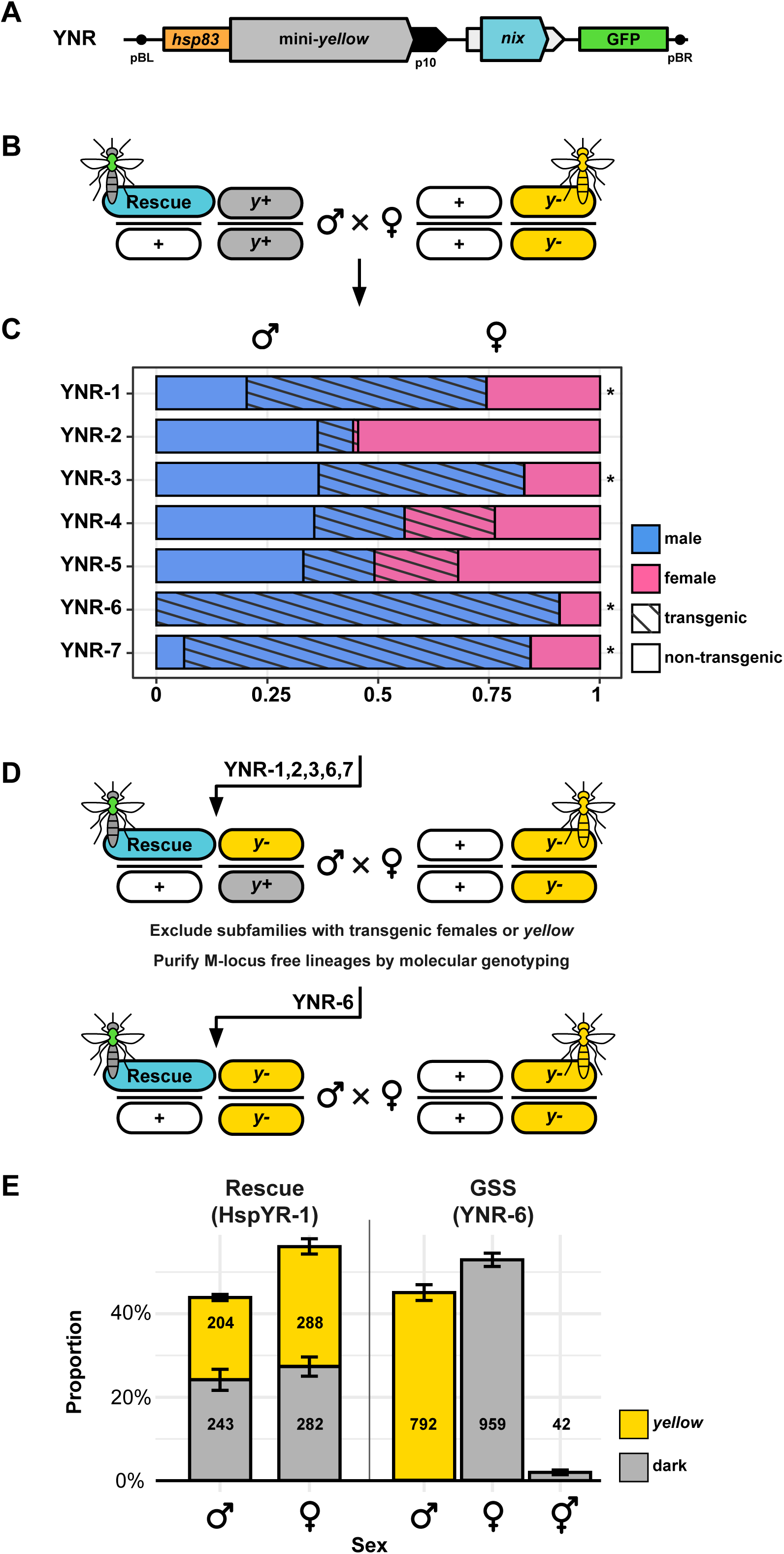
Combining mini-*yellow* rescue and sex conversion. **A)** Structure of the YNR construct created by adding *nix* isoforms 3 and 4 to the HspYR plasmid, which contains the *hsp83*-driven mini-*yellow* and the eGFP transformation marker. **B–D)** Crossing scheme for isolating functional integrations. **B)** Transgenic male isofounders are crossed with *yellow* females. **C)** F1 progeny were analyzed for sex and transgenic status. **D)** F2 transgenic males from isofamilies with significant sex bias (Z-test, p < 0.05 indicated by *) were individually crossed to *yellow* females to exclude subfamilies producing transgenic females or *yellow* individuals and to purify M-locus free male lineages by molecular genotyping for the endogenous *nix* and *yellow*. **E)** Sex conversion and *yellow* rescue in crosses between YNR and *yellow* females, using HspYR as a control. Frequency of pigmentation phenotypes (dark and *yellow*) separated by sex in progeny are shown with error bars representing standard error and numbers indicating the total individuals recorded per group.

YRN-6 produced exclusively dark transgenic males and *yellow* non-transgenic females **(Fig. 3C)**. In contrast, YRN-4 and YRN-5 exhibited high rates of transgenic females, suggesting failure of the transgene to masculinize genetic females. In families YRN-1,2,3,7 both transgenic and non-transgenic males were recovered, implying that their male founders were true males, transmitting both the M-linked *nix* and the YRN-linked *nix*. Despite multiple backcrossing rounds, these families did not yield pure-breeding pseudomales deprived of the endogenous M-locus.

We therefore focused on transgenic males from family YRN-6. Molecular genotyping confirmed that YRN-6 males lacked the endogenous M-locus (**Fig. S2**). To validate the function of YRN as a sexing system, we crossed *yellow* females either to YRN-6 pseudomales or HspYR-1 males that harbor only the *hsp83*-driven mini-*yellow*. In crosses involving HspYR-1, dark and *yellow* phenotypes segregated equally in male and female pupae (χ² ≈ 2.21, p = 0.137). In YRN-6, all female pupae recovered were *yellow*, while all phenotypically male pupae were dark (χ² ≈ 1747, p < 0.001) (**Fig. 3E**). A small proportion (4.2%) of pseudomales displayed intersex phenotypes, including intermediate genitalia, antennae and mouthparts. These intersexes were unable to blood-feed or mate. Based on these results we established YRN-6 as our first pure-breeding GSS strain that has now been maintained stably for more than 20 generations.

### Masculinization of gene expression in GSS pseudomales

To explore the effect of *nix-*based sex-conversion on sex-biased gene expression, we performed RNA-seq on adult whole bodies from WT males, WT females, GSS pseudomales, and GSS females. First, we assessed the correlation of genome-wide gene expression profiles, finding that GSS pseudomales closely resembled WT males (Spearman r_s_ =0.955 ± 0.001) despite being genetically identical to GSS females besides the insertion of the YRN construct **(Fig. 4A and Table <u>S1</u>)**. Next, we compared the expression of six well-known sex-biased genes - *nix, doublesex, fruitless, myo-fem, myo-sex*, and *yellow* **(Fig. 4B)**. For *nix* we found no reads mapping from either WT or GSS females, as expected. Both males and pseudomales on the other hand had good coverage over exons, but pseudomale coverage was limited to the two exons included in the *nix* transgene (isoform 3-4). Consistent with expression of *nix,* sex-specific splicing of the downstream sex-determination genes *doublesex* and *fruitless* was faithfully executed in pseudomales, mirroring the splicing in WT males and resulting in the skipping of female-specific exons. The female-specific flight muscle protein *myo-fem* was not expressed in pseudomales, but its male-specific paralog *myo-sex* was (O’Leary & Adelman 2020). Mapping to the *yellow* gene indicated that pseudomales express the *yellow* rescue at much higher levels than endogenous *yellow*, which is consistent with our use of the ubiquitous *hsp83* promoter. As expected, we observed a dip in mapped reads from GSS females in the position corresponding to the sgRNA target site, which likely is an outcome of indels contained in the *yellow* mutant alleles **(Fig. 1B)**. These results supported the conclusion of successful phenotypic masculinization and pigmentation encoded from the two rescue transgenes.

**Figure 4.**
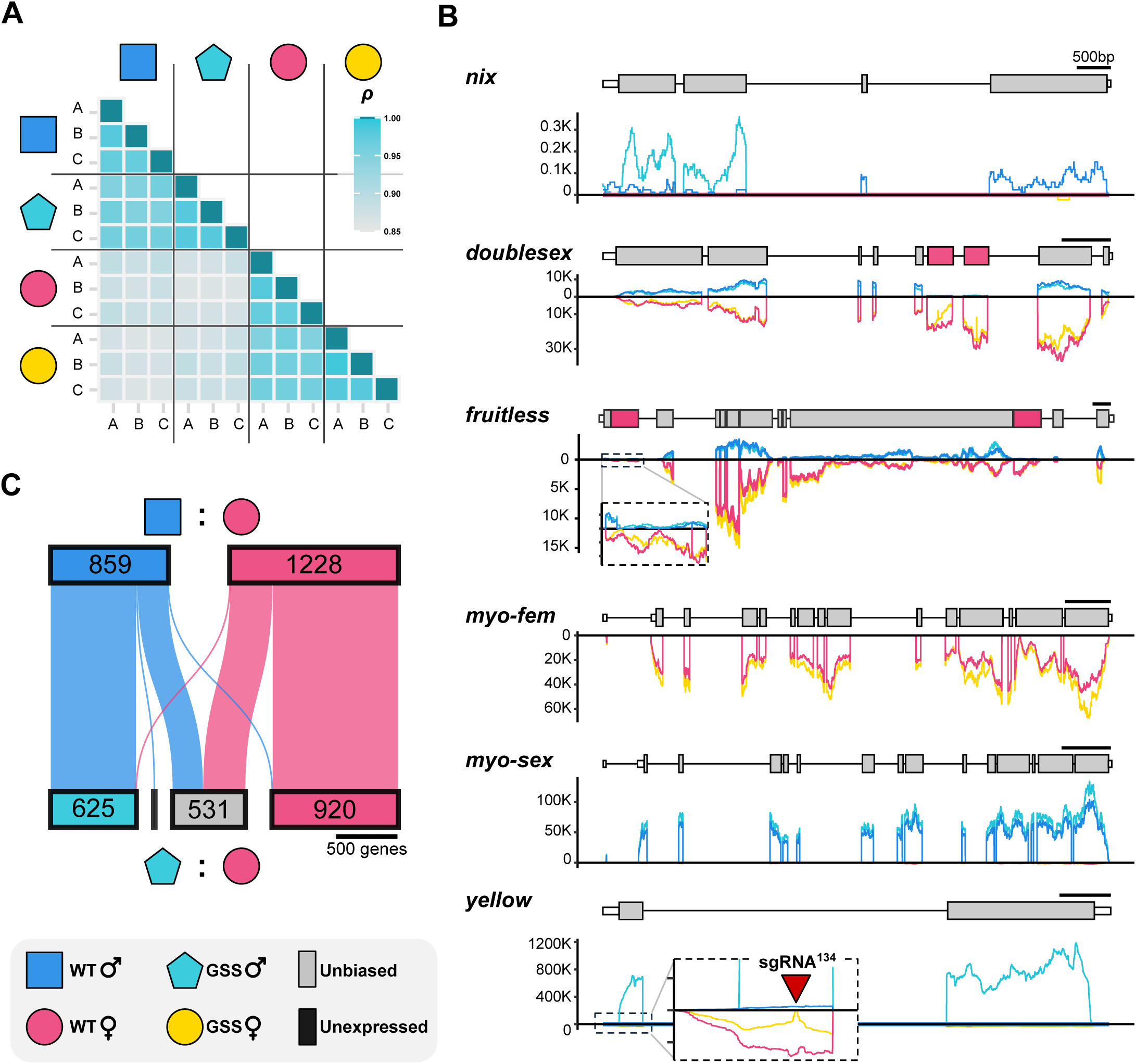
Sex-biased gene expression in sex converted GSS pseudomales. **A)** Spearman correlation matrix of genome-wide expression in sequenced samples. Higher correlation values indicate greater similarity in transcriptomic profiles. **B)** RNA-seq coverage plot for sex-linked markers (*myo-fem, myo-sex, yellow*) and key sex determination genes (*nix, doublesex, fruitless*). Exons are indicated in grey, except for female-specific exons that are indicated in pink. Introns not to scale. Coverage for male samples is shown on the positive Y-axis, while female coverage is reflected on the negative Y-axis for clarity. **C)** Differential expression analysis comparing WT females versus WT males (top) or GSS pseudomales (bottom), indicating changes or retention of sex-biased direction between analyses.

To catalog sex-biased gene expression, we ran differential expression analysis comparing WT males and females to compile a list of *Ae. albopictus* sex-associated markers, containing genes expressed in a sex-specific or significantly sex-biased manner. This catalog included 12.1% of the 17189 expressed genes, comprising 859 male-markers and 1228 female-markers. We then ran a second differential expression analysis comparing gene expression between GSS pseudomales and WT females, to track changes in sex-biased expression with WT females acting as a reference **(Fig. 4C)**. Overall, most markers retained their sex-bias - 623 (72.53%) and 917 (74.67%) for male- and female-markers, respectively. Levels of sex-biased expression (fold change) of markers that retained their sex-specificity was significantly higher than markers losing sex-biased expression (Tukey-Kramer HSD α = 0.05), indicating that retention of sex-bias was more likely for strongly sex-biased genes **(Fig. S3)**. The majority of markers that changed sex-bias classification in the pseudomale-female analysis became unbiased (222 male-markers and 309 female-markers **Fig. 4C**). Only two female-markers were more highly expressed in pseudomales, one of which was *yellow*. Eleven male-biased genes were completely absent in pseudomales, suggesting their M-locus linkage or in *trans* regulation by M-locus factors. To evaluate M-locus linkage we used the chromosome quotient method (Hall *et al*. 2013) and male-specific kmers (Carvalho & Clark 2013; Papathanos & Windbichler 2018) using male and female whole-genome sequencing data (Palatini *et al*. 2017). Five of the eleven genes were confirmed to be M-locus linked by PCR using male and female genomic DNA **(Table S1, Fig. S3)**.

### Benchmarking the *yellow*-GSS for sex sorting

To begin exploring the potential use of our color-based sex separation marker for *Ae. albopictus*, we quantified the pigmentation intensity of male and female pupae from GSS and WT strains to assess variation between individuals. To control for life-history factors like age, food availability, density etc. and to enable precise comparisons between WT and GSS individuals, larvae from each strain were co-reared in shared containers under high-density protocols, approximating conditions of mass-rearing (Balestrino *et al*. 2014). Image analysis of pupal pigmentation intensity confirmed that GSS female pupae were significantly paler than GSS pseudomales and all WT pupae (Tukey-Kramer HSD α = 0.05) (**Fig. 5A**). This pigmentation difference was so pronounced that optical sex separation of GSS pupae based on color and size, as observed by the naked eye, was faster and less error-prone than sex separation of the WT strain using a dissecting microscope (**Fig. S5 and S6**).

**Figure 5.**
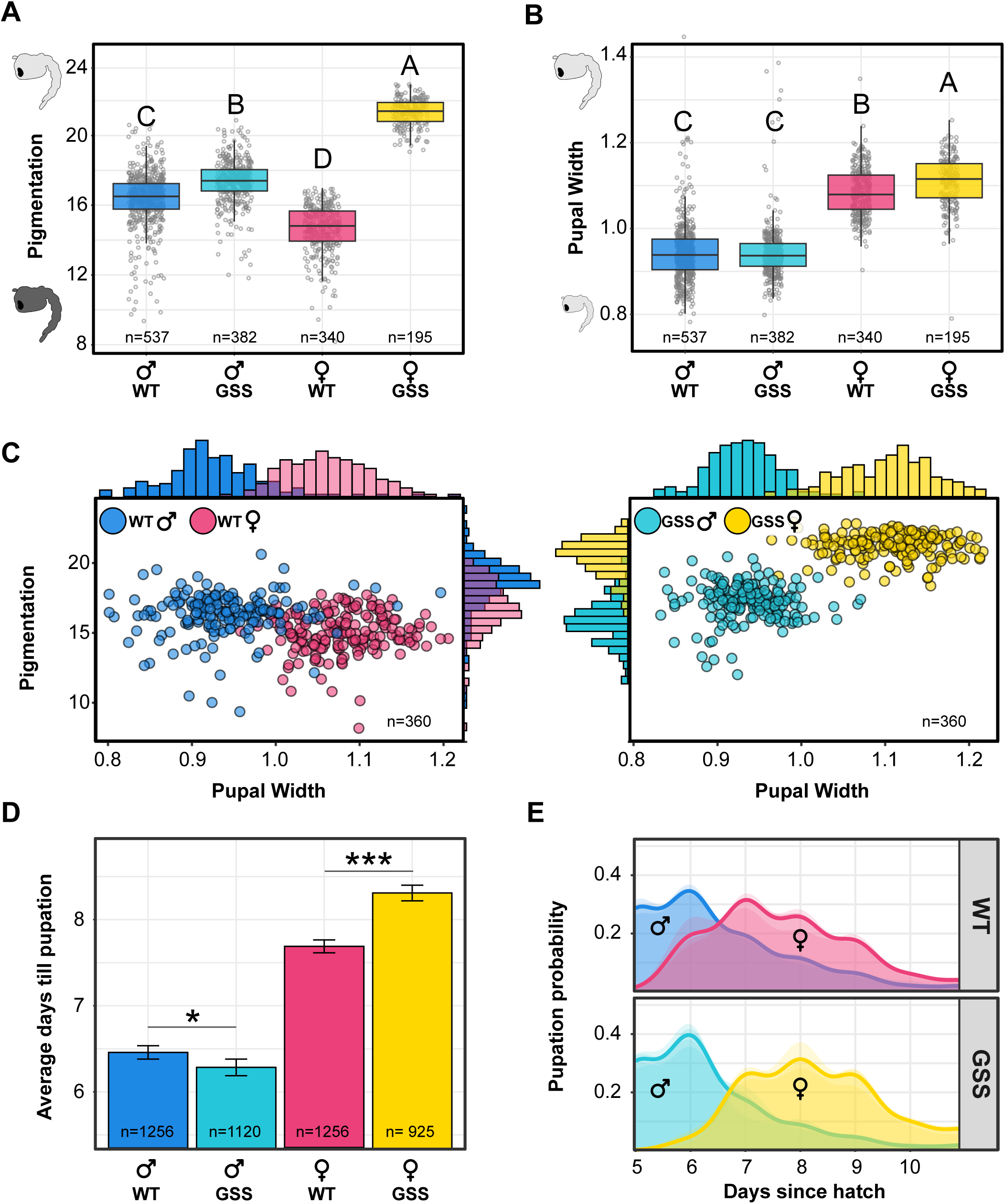
Engineered and enhanced sexual dimorphism of *yellow*-GSS. Measurements of **A)** Pupal pigmentation intensity (mean grey value) and **B)** pupal width (mm). Statistical significance for both metrics was determined using ANOVA followed by Tukey’s HSD test, with significant differences indicated by letter groups (p < 0.05). Error bars indicate standard error. **C)** Combined analysis of width and pigmentation levels. Each point represents an individual pupa, with histograms showing the distribution of phenotypes. **D)** Development time, measured as the average number of days until pupation. Statistical significance was determined by a student’s t-test (* for p < 0.05, *** for p < 0.001). Error bars indicate standard error. **E)** Pupation dynamics over time. Kernel density plots show daily pupation probability. Semi-transparent areas represent each of the four replicates in the experiment and solid lines indicate the cumulative probability.

We also collected data on pupal size and larval development dynamics for both strains, as proxies for mosquito fitness. We found no significant difference in the width, or any other size-related metric, of male pupae between the GSS and WT strain. Interestingly, GSS female pupae were slightly larger than female WT pupae (Tukey-Kramer HSD α = 0.05, **Table <u>S2</u>, Fig. 5B**). The intersection of size and color metrics enhanced the separation of males and females in the GSS strain compared to WTs (**Fig. 5C**). Insofar as insect size can reliably estimate mosquito fitness, these results suggest no detectable costs for GSS individuals compared to WTs. Importantly, they also suggest that this GSS is compatible with current size-based separation methods.

In terms of developmental dynamics, GSS pseudomales became pupae slightly faster than WT males and GSS females took significantly longer to pupate than WT females (**Table <u>S3</u>, Fig. 5D**). Combined, faster development of GSS pseudomales and slower development of GSS females increased the protandry index (accelerated male developmental time) to 27.8±0.82 in the GSS strain, compared to 17.4±0.29 in WTs. To test whether this enhanced protandry could be practically useful for operational use, we seeded triplicate WT and GSS high-density rearing trays, this time rearing each strain separately. Pupae were collected multiple times per day to improve the resolution of pupation dynamics **(Fig. 6A)**. Again, GSS females were the last to pupate, increasing the protandry index to 26.2±0.08 compared to 19.4±0.59 in WTs (ANOVA: F(3,7435)=1222, p<0.0001; Tukey-Kramer HSD: α=0.05; Fig. 6B). This approximate nine-hour delay in female pupation led to higher male recovery rates (proportion of recovered males from total males) for any rate of female contamination compared to the WT strain (**Table <u>S4</u>, Fig. 6C**).

**Figure 6.**
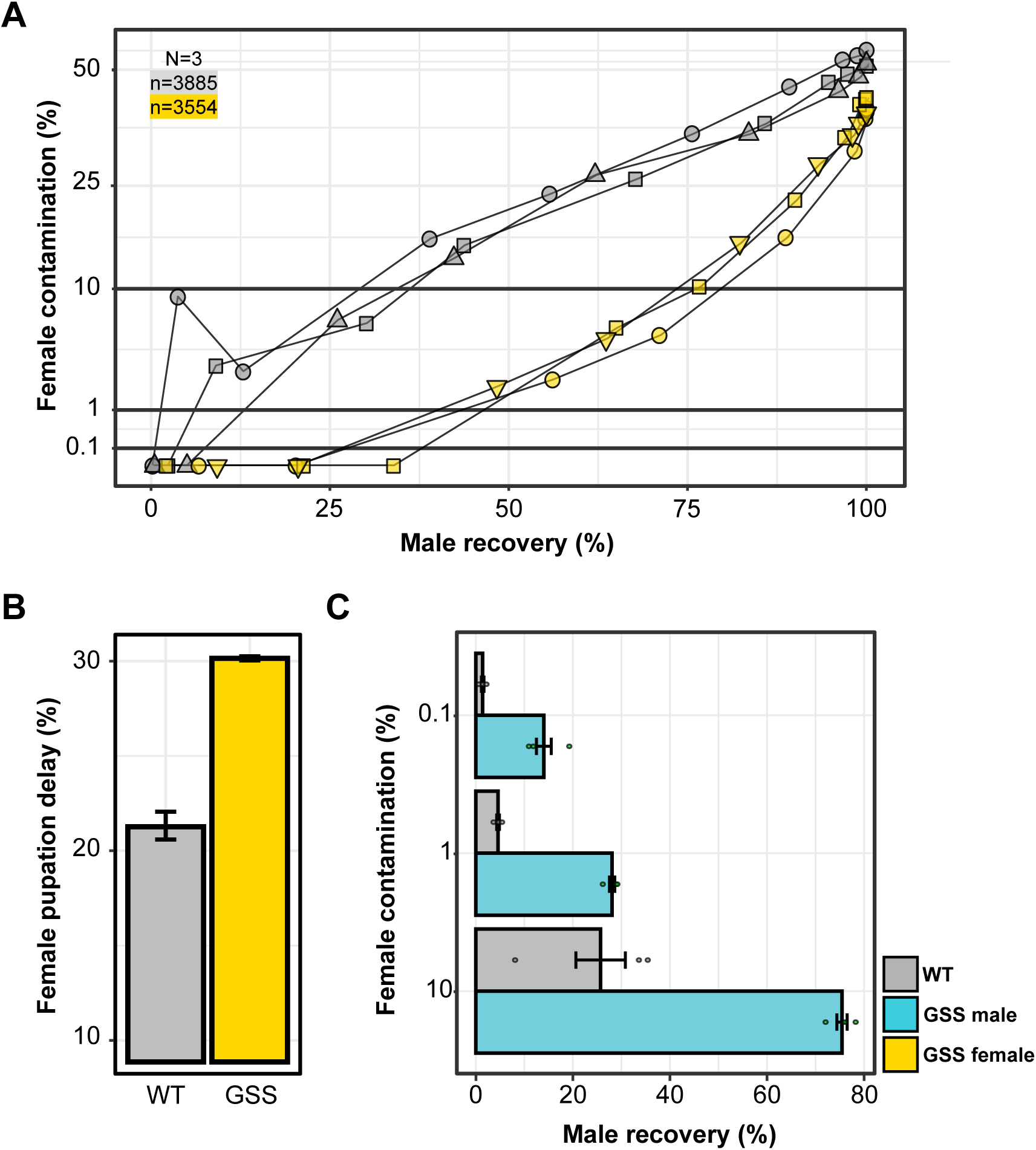
Enhanced protandry of *yellow-*GSS reduces female contamination. **A)** Pupation dynamics tracking male recovery and female contamination for the WT and GSS across three replicates for each line. **B)** Delay in the pupation time of females relative to males expressed as a percentage increase in development time. Error bars indicate standard error. **C)** Male recovery rates at three levels of female contamination comparing WT and GSS males.

The accelerated development of GSS pseudomales may reflect a trade-off between development speed and reproductive fitness (Pitnick *et al*. 1995; Yadav & Sharma 2014; Zeller & Koella 2016). To test whether their mating success was compromised, we assessed post-mating refractoriness in females first mated with either a pseudomale (eGFP/+) or a true male (dsRED/+), then exposed to males of the opposite type. Larval transgenic profiles were used to detect successful insemination by the second group of males. Such events could not be detected when females were mated in groups, but a few were detected in single-female mating assays. However, the likelihood of successful sequential insemination did not differ significantly between the two male types (z = -0.589, df = 38, p = 0.556). We concluded that females mated with GSS pseudomales were equally resilient to further mating attempts, as those mated with WT males (**Table <u>S5</u>**).

### Desiccation-sensitivity of GSS eggs

During the establishment of stable *yellow* and GSS populations, we observed considerable variability in the hatching rates of eggs laid by *yellow* females. Previous studies have shown that following oviposition the chorion of *Ae. albopictus* eggs undergo melanization via a process mediated by several *yellow*-family genes, which hardens the eggshell and confers desiccation resistance (Noh *et al*. 2020, 2021). Considering that *yellow* eggs did not darken like WT eggs **(Fig. 1G)** and were too fragile for microinjection, we hypothesized that the absence of maternal *yellow* could compromise desiccation resistance and/or embryonic dormancy - traits that have enabled the long-distance dispersal of this mosquito as desiccated, dormant eggs (Girard *et al*. 2024; Swan *et al*. 2022).

To investigate this, we collected twelve egg bowls from cages containing *yellow* males crossed to WT and GSS females and recorded for each egg bowl, the total number of eggs laid by WT (dark eggs) and GSS (yellow eggs) females. During storage, we randomly selected three egg batches every seven days for hatching and screened hatched larvae for cuticle color to calculate relative hatching rates. Overall, GSS eggs exhibited significantly lower hatching than WT eggs at all time points (paired t-test, t(11)=10.6, p < 0.001, ANCOVA phenotype: F(1,18) = 10.97, p = 0.0039), storage duration: (F(1,18) = 9.71, p = 0.0060), Tukey-Kramer HSD α = 0.05) **(Fig. 7A)**. While the viability of WT eggs began to decline only after the third week of storage—remaining above 50% until the end of the experiment—GSS eggs were completely unviable by the fourth week **(Fig. 7A)**. To test whether the increased mortality of GSS eggs was due to elevated desiccation sensitivity, we transferred fully-matured, 4-day-old GSS and WT eggs from moist filter papers to glass slides and monitored their morphology as the surrounding environment dried via evaporation under the microscope. Within a few minutes, most *yellow* eggs displayed visible signs of desiccation, ultimately resulting in the irreversible collapse of their chorionic structure rendering them nonviable. Conversely, WT eggs retained their structural integrity and remained unaffected **(Fig. 7B-C)**. These findings show that the *yellow* mutant background of GSS females imparts significant desiccation sensitivity that is maternally-defined, a trait that could serve as a useful barrier to prevent gene flow or establishment of incompatible symbionts in field population from the unintentional, rare release of contaminating females. Importantly, since identifying the desiccation sensitivity of the GSS eggs, we modified our protocols for egg storage to maintain optimal moisture and to hatch stored eggs within two weeks from oviposition - practices that ensure a healthy and stable maintenance of the GSS population in our insectary for the last one and half years.

**Figure 7.**
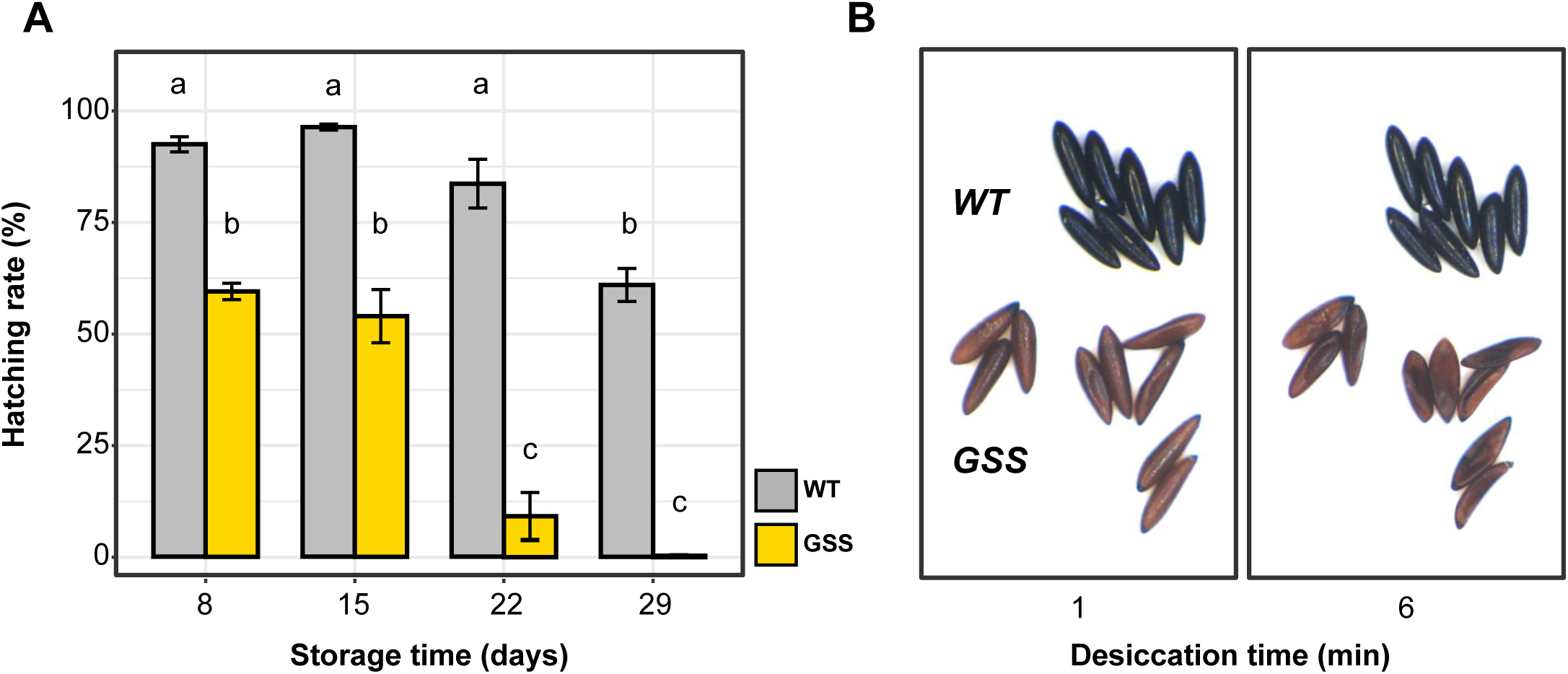
Desiccation sensitivity of GSS eggs. **A)** Hatching rates of stored WT and GSS eggs tracked over one month. Hatching rates were calculated as the proportion of eggs that successfully developed into L2 larvae. Error bars denote standard error. Statistical significance was determined by ANCOVA followed by post-hoc Tukey-Kramer HSD, with significant differences indicated by letter groups (α = 0.05). **B)** Time-lapse images of mature, 4-day-old eggs WT and GSS cages mounted on a dry glass slide and photographed 1- and 6-minutes following removal from moist egg papers.

## Discussion

The VIENNA-8 GSS of the Mediterranean fruit fly (*Ceratitis capitata*) has for decades now served as the gold standard for insect sex separation, providing a reliable and scalable solution for medfly SIT programs (Augustinos *et al*. 2017; Franz 2005). Despite extensive efforts and coordinated research to develop similar GSSs in other insect species of medical and agricultural importance, progress has been limited, primarily due to the unpredictability of classical genetic approaches. As a result, genetic control programs for *Aedes* mosquitoes currently rely on naturally occurring sexual dimorphisms for female elimination (Crawford *et al*. 2020; Lutrat *et al*. 2019; Mamai *et al*. 2024).

The most widely used method is size-based sorting at the pupal stage, where female pupae are slightly larger than males. However, this approach presents several challenges: 1) size dimorphism is a continuous trait with significant overlap between the sexes leading to female contamination (Wormington & Juliano 2014); 2) sexing at the pupal stage requires rearing both sexes until pupation resulting in wasted resources; and 3) manual size-based sorting is labor-intensive and difficult to scale for mass-production. While recent advances in automated sorting technologies, such as robotic systems leveraging size or other morphological differences, may help address some of these limitations, they remain costly and complex to implement (Crawford *et al*. 2020; Lutrat *et al*. 2023; Mamai *et al*. 2024). In contrast, a well-designed GSSs could offer a more practical, cost-effective and scalable solution for mosquito sex separation (Papathanos *et al*. 2018).

In this study we introduce a novel and versatile platform for the development of GSSs in the important arboviral vector *Ae. albopictus*. Our platform is based on the CRISPR-mediated disruption of native mosquito genes encoding selectable phenotypes, followed by their genetic rescue using sex-converting transgenes. As a proof-of-concept, we selected the highly-conserved *yellow* gene - one of the first phenotypic mutants discovered in the fly room of Thomas Hunt Morgan (Morgan 1911). We selected this gene for its relatively simple structure and lack of multiple isoforms. Based on previous work in *Ae. albopictus*, homozygous null mutants were expected to display a clearly visible phenotype throughout development, and to be both viable and fertile (Liu *et al*. 2018), which we confirmed by generating a mutant strain using sgRNA^134^ targeting the first exon and established a stable homozygous strain.

To genetically rescue cuticle pigmentation, we generated constructs containing the spliced *yellow* coding sequence that we called mini-*yellow*. The EnYR construct containing mini-*yellow* flanked by its native regulatory regions failed to complement pigmentation in all transgenic strains generated, suggesting that the selected regions lack critical promoter and/or enhancer regions. A second construct, HspYR, expressing mini-*yellow* from validated regulatory regions of the *Ae. albopictus hsp83* gene successfully restored pigmentation in the *yellow* mutant background. RNAseq expression data revealed that the *hsp83* regulatory regions drove high levels of expression of the mini-*yellow* in adults, suggesting that this promoter could be useful for the development of other GSSs with selectable traits requiring ubiquitous expression.

Building on recent work (Lutrat *et al*. 2022; Zhao *et al*. 2022), which demonstrated that *nix* alone is sufficient to masculine genetic females into fertile males, we incorporated *nix* into our mini-*yellow* construct, effectively coupling dominant pigmentation with masculinizing sex conversion. Among the seven independent strains generated, YRN6 was the only one in which all phenotypically dark males carried the transgene, confirming exclusive linkage between male sex and the *nix* transgene. In contrast, three other strains produced both transgenic and non-transgenic males, indicating that their founders were likely genetic males rather than sex-converted pseudomales. Attempts to isolate males lacking the native M-locus from these strains, by individually crossing fluorescent males to *yellow* females, failed to recover any family containing exclusively sex-converted males, suggesting incorrect expression of the *nix* transgene from these integrations and lack of pseudomale fertility in these strains.

Overall, our results confirm that *nix*-based sex conversion offers a practical alternative to other methods for tightly-linking rescue constructs to maleness, overcoming key technical challenges associated with reciprocal translocations and Y-chromosome knock-ins. Reciprocal translocations impose significant male fertility costs due to gamete aneuploidy (Benet *et al*. 2005). Y-chromosome knock-ins, on the other hand, remain technically challenging due to the highly repetitive nature of male-specific chromosomes, making precise insertions difficult. To date, successful Y-chromosome knock-ins have only been achieved in *D. melanogaster,* despite significant efforts - both by our group and others - to adapt this approach for other insect species (Buchman & Akbari 2019; Gamez *et al*. 2021). Additionally, even when Y-linked insertions are obtained, integrations frequently suffer from reduced or variegated transgene expression, likely due to the heterochromatic environment of the Y chromosome (Berloco *et al*. 2014; Dimitri & Pisano 1989; Gamez *et al*. 2021). In *A. albopictus*, *nix*-based sex conversion bypasses these limitations by enabling a stable, autosomal approach to masculinization, providing a more reliable and scalable solution for genetic sexing.

Most insect species are not as malleable as *Ae. albopictus* in manipulations of their sex determination pathway. Even when master sex determination genes are known, their expression in genetic females can be insufficient to fully convert them into fertile males, for example by requiring functions of other male-specific genes present in Y-chromosomes or M-loci (Aryan *et al*. 2020). Alternatively, transgenic expression of primary male-determiners can be lethal, for example by interfering with the dosage compensation pathway in genetic females and resulting in misexpression of X-linked genes.(Criscione *et al*. 2016; Krzywinska & Krzywinski 2018; Qi *et al*. 2019). Our transcriptomic analysis confirmed that despite their genetic female background, pseudomales exhibited a gene expression profile highly similar to that of WT males. This included the correct splicing of well-known sex-determination genes like *dsx* and *fru (Salvemini et al. 2011, 2013)* but also most other sex-biased genes, which we cataloged here for this first time for this mosquito species. Among the male-markers we found an interesting subset of eleven genes, whose expression was not detected in GSS pseudomales suggesting M-locus linkage, or their transcriptional regulation by M-locus linked factors other than *nix*. We confirmed the localization of five of these within the M-locus, of which one (LOC134287234) is predicted as non-coding and four as protein coding genes. LOC109432934 contains a leucine-rich repeat domain, LOC134285312 a sperm tail domain, LOC134289344 a YqaJ-like viral recombinase domain and LOC134285028 lacks any known motifs.

In terms of sex separation, we benchmarked the fitness and performance of the *yellow*-GSS, measuring differences in pigmentation intensity between male and female pupae. Our results confirmed that GSS females were significantly paler than GSS pseudomales, making sex sorting based on color highly efficient. The pronounced contrast in pigmentation allowed for rapid and more reliable visual sorting compared to the traditional method of pupal sexing under a dissecting microscope or based on size. This suggests that *yellow*-GSS provides a robust improvement to current sorting methods, providing a clear, visible indicator of female contamination. Color based sorting could be useful as a standalone trait for imaging- and algorithm-assisted automated sorting, or could be used as a secondary marker in future designs involving high-throughput traits like temperature-sensitive lethals.

Beyond pigmentation, we also examined pupal size and larval development dynamics to evaluate potential fitness costs associated with the GSS genotypes. Pseudomale pupal size and development was comparable to WT males. GSS female pupae were slightly larger than WT females, a feature that enhanced sex separation accuracy, when combined with pigmentation differences. Interestingly, homozygosity for *yellow* in GSS females led to slower larval development, significantly increasing protandry—the earlier emergence of males relative to females. In WT *Ae. albopictus*, protandry alone is usually insufficient for reliable sex separation due to overlapping male and female development times (Bellini *et al*. 2018). However, the enhanced protandry observed in the GSS expanded the temporal gap between male and female pupation, improving sorting efficiency and reducing female contamination. This aligns with findings by Bellini et al. (Bellini *et al*. 2018), who noted that while natural protandry exists, it is not pronounced enough for effective sex separation. By amplifying this developmental difference, *yellow*-GSS offers an additional trait that could have practical applications for mosquito control programs.

To evaluate the suitability of our GSS pseudomales for genetic control programs, we assessed their ability to induce post-mating female monogamy, also known as mating refractoriness (Sutter *et al*. 2021). We found that pseudomales induced levels of female post-copulatory refractoriness comparable to those of true males, supporting their potential utility in genetic control strategies. Male mating competitiveness can generally be divided into two categories: pre-mating and post-mating competitiveness. Pre-mating competitiveness refers to the ability of released males to successfully mate with females in competition with wild males, reflecting factors such as courtship vigor and attractiveness. Post-mating competitiveness, in contrast, refers to the ability of released males to successfully fertilize eggs and induce refractoriness in females, preventing or significantly reducing subsequent matings with wild males. Post-mating competitiveness is particularly critical for genetic control programs targeting insect species whose females naturally exhibit strong mating refractoriness under field conditions, as is the case with many mosquitoes (Alphey et al. 2010; Knipling 1955; Lance & McInnis 2021; Whitten & Mahon 2005). Even if released males exhibit normal pre-mating competitiveness, population suppression efforts will fail if these males do not effectively induce refractoriness. While suboptimal pre-mating competitiveness can typically be compensated for by increasing male release ratios, low post-mating competitiveness cannot be easily mitigated. Previous studies evaluating *nix*-masculinized pseudomales reported reduced mating competitiveness in laboratory cages using 1:1 ratios with true males (Lutrat et al. 2022; Zhao et al. 2022). In these experiments, competitiveness was measured as the fraction of offspring sired by pseudomales compared to true males, a variant of Fried’s index typically used for measuring sterile male competitiveness. However, it is now generally well established that such measurements of mating competitiveness can result in high levels of variation in the calculated competitiveness from each cage with competitiveness changing according to the male release ratio even though it theoretically should not, and that small-cage experiments promote the otherwise rare occurrence of multiple insemination events (Yamada et al. 2024). Therefore, reliable evaluations of GSS pseudomale fitness will require additional testing including multiple release ratios and ideally larger, semi-field conditions, which were outside the scope of this study. Our results demonstrate that GSS pseudomales possess post-mating competitiveness comparable to true males, justifying further investigations into their overall competitiveness and fitness. Future studies should explicitly assess pre- and post-mating competitiveness separately across a range of ecologically relevant release ratios and evaluate their interactions with genetic control technologies, such as sterilization, in conditions more representative of natural populations.

To evaluate the compatibility of our GSS pseudomales for genetic control programs, we assessed their ability to induce post-mating female monogamy, also called mating refractoriness. We found that pseudomales induced levels of post-copulatory refractoriness in females comparable to those of true males. Generally, there are two types of male mating competitiveness that can be evaluated for new strains developed for genetic control: premating and postmating competitiveness. Premating competitiveness refers to the ability of a released male to successfully mate with females in competition with wild males. Post-mating competitiveness refers to the ability of a released male to successfully fertilize eggs and prevent or reduce female remating after an initial mating event, if the females of the insect targeted become refractory to further mating (i.e. are monogamous) in the field.

In genetic control programs, post-mating competitiveness is especially critical (Alphey et al. 2010; Knipling 1955; Lance & McInnis 2021; Whitten & Mahon 2005), because even if released males mate effectively (normal pre-mating competitiveness), population suppression will fail if these males are not able to induce refractoriness in mated females (suboptimal post-mating competitiveness). Furthermore, low pre-mating competitiveness, while a disadvantage, can be mitigated through male releases at higher release ratios, but low post-mating competitiveness cannot be easily compensated. As a result, poor post-mating competitiveness would be a significant limitation, as it directly impacts reproductive outcomes and cannot be overcome with higher release ratios. Previous studies by Lutrat and Zhao (Lutrat *et al*. 2022; Zhao *et al*. 2022) found that pseudomales competitiveness was reduced in small-cage, 1:1 competitiveness assays against true males (containing the M-locus). In these studies, competitiveness was measured as the fraction of pseudomale offspring produced by females exposed to pseudomale and true males - effectively a variation of how sterile male competitiveness is measured known as the Fried’s Competitiveness index, which counts hatching rates and normalized for release ratios (Fried 1971; Hooper & Horton 1981). However, it is now well established that measurements of mating competitiveness based on this design, especially when using standard laboratory cages, can result in significant variation in the computed competitiveness and to reliably measure true competitiveness multiple release ratios need to be tested and ideally use large semi-field cages (Bond *et al*. 2021; Chen *et al*. 2025). Importantly, our first results on GSS pseudomale suggest that post-mating competitiveness is similar to true males, offering sufficient justification for the continued evaluation of the competitiveness and fitness of GSS pseudomales. Future studies should separately assess both pre- and post-mating competitiveness across a range of release ratios, and consider their interaction with the implemented genetic control technology like sterilization in an ecological meaningful context.

A notable characteristic of the *yellow* mutant is the desiccation sensitivity of GSS eggs. The failure of eggs laid by GSS females to melanize properly resulted in a marked reduction in their viability over time. While this trait posed an initial challenge that required adjustments to rearing protocols, simple measures such as maintaining optimal humidity of egg papers and hatching stored eggs within two weeks have ensured stable colony maintenance in the laboratory. We anticipate that these protocols could be easily adapted for future mass-rearing operations, ensuring the strain’s scalability (Zheng *et al*. 2015). In the context of field deployment, this trait could serve as an inherent containment mechanism, minimizing the risk of accidental female releases and reducing the potential for unintended gene flow or *Wolbachia* establishment in target populations.

Looking ahead, our findings suggest that color-based sex sorting can be both faster and more accurate than pupal size sorting alone. The *yellow* mutant, besides being an excellent color marker, also offers useful secondary phenotypes like slow larval development and egg desiccation sensitivity. Future studies could focus on developing high-throughput sorting technologies to fully exploit the advantages of this strain. While the *yellow* phenotype offers a strong standalone visual marker, future iterations of this GSS platform could integrate next-generation selectable traits, such as temperature-sensitive lethals, to enhance scalability further. A key strength of our GSS development framework is its adaptability—new selectable marker mutants and their corresponding rescues can be iteratively introduced and expanded, enabling the system to evolve alongside future advancements in genetic control technologies.

## Methods

### Mosquito strains and rearing

The FPA (Foshan) *Ae. albopictus* strain was used as the laboratory WT strain in all experiments. All strains were reared at 27±1°C, 75±5% humidity with 14:10 (LD) photoperiod regime. Larvae were reared in 160 x 135 mm plastic containers in 300ml demineralized water at a density of up to 1.33 larvae per milliliter. The rearing tank was provided with a daily supplement of fish-food (‘Essence’, Alltech-coppens,Leende, NL). Pupae were manually collected every 48 hours, and transferred to the designated cages. Adult mosquitoes were maintained in cages and given access to TGS solution (10% sucrose, 0.1% methyl paraben). Female blood meals consisted of heparinized bovine blood augmented with 5mg/ml ATP. Blood meals were provided using Hemotek feeding system (Discovery Workshops, Accrington, UK). Four days after blood feeding, an egg-bowl was introduced into each blood-fed cage. After 24 hours, egg-bowls were removed and egg papers were transferred to Petri dishes. Petri dishes were left to dehydrate for 24 hours before being parafilm sealed to maintain some moisture for long-term storage. To synchronize hatching, mature stored eggs (≥96 hours old) were immersed in deoxygenated hatching solution consisting of 0.25 mg nutrient broth (Merck-Sigma Aldrich, USA) and 0.05 mg baking yeast dissolved in 750 ml of distilled water and incubated overnight at room temperature. Hatched larvae were collected after 6 hours as L1, or the following day as L2 and moved to rearing containers.

### Generating a *yellow* mutant strain

We selected sgRNA-134 (CACGTAGTCCCCACTGGCCA-GGG) to target the first exon of the *yellow* gene (LOC109402083 in AlboF5) using CRISPOR (Concordet & Haeussler 2018). Ribonucleoprotein injection mixes containing SpCas9 (300 ng/μL) and sgRNA (80 ng/μL) (IDT, Coralville, IA) were pre-incubated at 37°C for 30 minutes and microinjected into syncytial blastoderm embryos following standard protocols (Fuchs *et al*. 2013). Mosaic individuals were backcrossed to WT mosquitoes to establish isofamilies. To minimize fitness costs and account for the proximity of *yellow* to the non-recombining region of chromosome 1 (which contains the M and m loci), heterozygotes from different families were crossed to establish a homozygous *yellow* line. DNA from a mixed-sex pool of F5 *yellow* individuals was extracted, and the target site was PCR amplified and sequenced using next-generation sequencing (NGS; Hy-labs, Israel). Mutant alleles were characterized and quantified, and predicted protein structures were generated using AlphaFold (Jumper *et al*. 2021).

### Plasmid construction and transgenesis

For the EnYR mini-*yellow* rescue, the *yellow* coding sequence was amplified in two fragments: exon 1 (with 1,100 bp upstream regulatory region) and exon 2 (660 bp), omitting the large intron. These fragments were assembled using Gibson assembly into an Opie2:eGFP piggyBac transformation plasmid. For the HspYR plasmid, a 2076 bp fragment of the *hsp83* promoter (LOC109407918) was cloned upstream of the mini-*yellow* coding sequence followed by the P10 terminator (Li *et al*. 2017); Addgene plasmid #100580). To generate the YNR plasmid, the *nix*3&4 isoform (Lutrat *et al*. 2022); Addgene plasmid #173667) was inserted into the HspYR plasmid. Cloning was performed using HiFi DNA Assembly (New England Biolabs) and Q5 Hot Start High-Fidelity PCR. Plasmids were validated by sequencing and microinjected at 200 ng/μL along with 40 ng/μL of a mosquito codon-optimized piggyBac transposase helper plasmid (Lutrat *et al*. 2022). Individuals transiently expressing eGFP were backcrossed to WT to establish transgenic isofamilies.

### Genotyping for endogenous and transgenic components

Specific primer sets were designed to distinguish between endogenous and exogenous components (**Table <u>S6</u>**, **Fig. S2A**). For the *yellow* gene, forward primers targeting the 5′ UTR and *hsp83* promoter were paired with reverse primers matching the unaltered sgRNA134 cut site and *yellow* exon 2. To differentiate between endogenous and transgenic *nix*, a triplex PCR was employed using a forward primer for exon 1 and two reverse primers, one targeting an endogenous intronic gap and another specific to the transgenic SV40 terminator. These assays were used to genotype DNA from transgenic dark male isofounders and to identify lineages deprived of endogenous elements (**Fig S2B)**.

### Testing *yellow* rescue from transgenic constructs

Ten transgenic males carrying either the YRN construct or the non-masculinizing HspYR construct in the WT genetic background were crossed to ten *yellow* females. Two replicate cages were used for each cross, with each cage undergoing three rounds of blood feeding and egg laying (technical replicates). The sex, phenotype, and transgenic profile of pupae from each gonotrophic cycle were recorded to calculate sex-specific phenotype distributions.

### Sex-biased gene expression analysis

Four replicate samples, each comprising three 4-day-old flying males or females from the WT and GSS strains, were collected. Anesthetized mosquitoes were transferred to tubes containing 2.3 mm zirconium-silicate beads (BioSpec Products, USA) and ice-cold TRI-reagent (Zymo Research Corp., USA). Whole-body RNA was extracted using a Minilys bead-beater (Bertin Corp., USA), followed by two chloroform washes, ethanol suspension, and purification on a Zymo-spin IICR column (Zymo Research Corp., USA). mRNA libraries were prepared using poly-A selection and sequenced on an Illumina NovaSeq 6000 with 100 bp paired-end reads, yielding approximately 45 million reads per sample. Sequencing data are available in NCBI under bioproject accession PRJNA1100762. Adapter sequences and poly-A/T stretches were removed using cutadapt (Martin 2011), and reads shorter than 30 bp were discarded. Remaining reads were aligned to the *Ae. albopictus* reference genome (AalbF5) using the STAR aligner (Dobin & Gingeras 2015) in EndToEnd mode with an outFilterMismatchNoverLmax parameter of 0.04. Duplicate reads were removed using PICARD MarkDuplicates (“Broad Institute” 2025), and gene expression was quantified with htseq-count (Putri *et al*. 2022) using the AalbF5 GTF annotation. Principal component analysis (PCA) was used to select the three most similar samples per group for differential expression analysis with DESeq2 (Love *et al*. 2014), filtering for a log₂ fold change >2 and adjusted p-value <0.05. Genes were classified as specifically expressed if they were detected in all samples of one group (with a cumulative count ≥5) and absent in the other.

### Analysis of M-locus linkage of male-associated genes

To confirm the linkage to the M-locus of genes expressed in WT males but not in pseudomales, two computational approaches were used: the chromosome quotient (CQ) method (Hall *et al*. 2013) and male-specific kmers (Carvalho & Clark 2013; Papathanos & Windbichler 2018) (**Table S1**). CQ values were calculated by mapping male and female whole-genome sequencing reads (Palatini *et al*. 2017) to transcripts and coding sequences in the AalbF5 assembly (GCF_035046485.1) using bowtie with parameters -v0, -a, and --suppress 1,2,4,5,6,7,8,9 (Langmead *et al*. 2009). The CQ for each sequence was determined as the ratio of female to male normalized reads. Additionally, 25 bp kmers were generated from each sex-specific library using jellyfish (Marçais & Kingsford 2011), kmers present at least ten times in male libraries but absent in female libraries were identified as male-specific. These kmers were mapped to putative M-linked sequences using bowtie, and the total length covered by male-specific kmers was measured. Finally, PCR on genomic DNA from pools of 20 males or females was used to validate M-locus linkage predictions.

### Quantifying pupal color, size and protandry

Images of pupae in water-filled petri-dishes were captured using a Nikon D7100 camera with a Laowa 60 mm f/2.8 2X Ultra-Macro lens mounted on a tripod. Image analysis was performed with ImageJ. Size measurements were calibrated using the petri-dish diameter as a reference. The script for region-of-interest (ROI) identification and parameter measurement is provided in Supplementary File <u>S1</u>, along with sample images of WT and GSS pupae in the (**Fig. S5**).

Synchronized larvae were obtained by simultaneously immersing WT and GSS eggs in hatching solution. L1 larvae from each strain were counted, and three replicates of 1,500 larvae per strain were seeded into 500 mL of water (3 larvae/mL) in trays with a 325 × 175 mm surface area (2.6 larvae/cm²). Larvae were fed daily with TetraBits Complete fish food (Tetra, Melle, Germany), and trays were monitored for the onset of pupation. Once formed, and over a five-day period, pupae were collected at 10-time intervals, manually sexed, and counted. Recovery rates for males and females were calculated based on the total number collected from each tank. Protandry was calculated for each replicate using an adaptation of the SBM (sexual bimaturism) index calculation following (Blanckenhorn *et al*. 2007) using the mean pupation time of females (F) and males (M) with the following formula : SBM= 100 x [2(F-M)]/(F+M)

### Mating refractoriness assays

To evaluate whether GSS pseudomales successfully induce post-mating refractoriness in females, we conducted two reciprocal mating assays. In the first experiment a group of 50 unmated, 5-day-old females were introduced to a cage containing either 25 GSS pseudomales (fluorescently labeled with eGFP) or 25 true males carrying a fluorescent marker (OpIE2:dsRED). Two hours after introduction, females were blood-fed and transferred to a cage containing 25 males of the opposite type for the remainder of the experiment. Four days later, three egg bowls were collected from each cage and the paternal origin of L2 larvae was determined using fluorescent screening. In the second experiment, in which single female insemination status was assessed, 25 females were individually introduced into a cage containing either 25 GSS pseudomales or transgenic true males. Each female was monitored from the moment of introduction for interception and mating. Following at least one successful copulation (lasting at least 20 seconds) the mated female was transferred to a cage containing 25 males of the opposite type for the remainder of the experiment. The cages were blood-fed on the same day, and females were individually collected four days later for egg laying. Single-female clutch analysis based on the L2 transgenic profile was subsequently performed for 20 females from each group.

### Evaluating desiccation-sensitivity of GSS and WT eggs

Six egg bowls were placed in a blood-fed cage containing an equal number (50 each) of WT and GSS females, both mated to *yellow* males. Prior to storage, the egg papers were divided in half, and the total number of eggs was documented on the 12 egg papers, distinguishing between *yellow* and WT eggs. Over the course of one month, three stored egg papers were randomly hatched each week. Larval phenotypes were evaluated three days post-hatching at the L2–3 stage, and hatching rates were calculated based on the total number of recovered larvae per strain.

### Statistical analyses

All statistical analyses were performed using R version 4.4.1 (2024-06-14 ucrt). Data were initially assessed for normality with the Shapiro–Wilk test and for homogeneity of variance using Levene’s test. For normally distributed data, comparisons between groups were conducted using one-way ANOVA—or ANCOVA when groups were compared across different time points, followed by Tukey’s HSD post hoc test for multiple comparisons. Two-level group comparisons were performed using either a T-test or a paired T-test when appropriate. Non-normally distributed data were analyzed with the Kruskal–Wallis test followed by Dunn’s post hoc test, and population phenotype proportions were compared using Chi-square and Z-tests. Statistical significance was set at α = 0.05 for all tests. Plots were created using the ggplot2 and networkD3 packages.

## Supporting information

Table S6 - primers list

## Acknowledgements

We would like to thank Dor Perets, Guy Ostrovsky, Shira Kehat, Gleb Ens and Ezra Bohbot for technical assistance. Special thanks to Kostas Bourtzis for his invaluable insights, longstanding support and guidance, which have greatly contributed to this work. We thank Mariangela Bonizzoni and Francesca Scolari for the Foshan strain. We thank Amir Szitenberg of the Mantoux Bioinformatics Institute of the Nancy and Stephen Grand Israel National Center for Personalized Medicine, Weizmann Institute of Science for technical support in the RNAseq analysis. This study benefited from discussions at meetings for the Coordinated Research Project D44003 on the “Generic approach for the development of genetic sexing strains for SIT applications”, funded by the International Atomic Energy Agency (IAEA).

## Ethics

All animals were handled in accordance and under the supervision of the ARO Institutional Animal Care and Use Committee approval number 2307-118-2-VOL-IL. All insect work was performed in facilities maintaining Arthropod Containment Containment Level 2. This work received Institutional Approval and relevant authorizations from the Israel Ministry of Environmental Protection and Ministry of Agriculture (#31/2019).

