## supplementary figures for "Mosquito Sex Separation using Complementation of Selectable Traits and Engineered Neo-Sex Chromosomes"

### Supplementary Figure S1

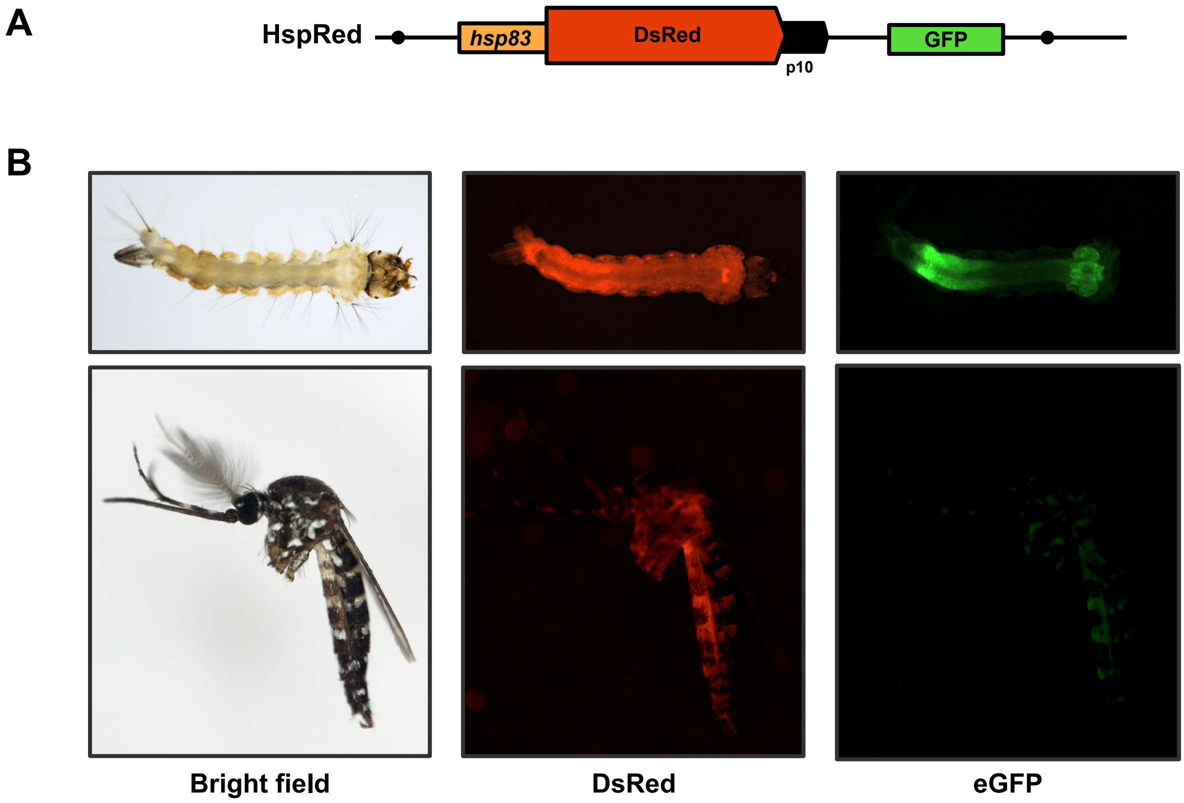


**Supplementary Figure 1. Validation of *hsp83* regulatory regions using a fluorescent reporter. A)** Structure of Hsp83Red construct containing the *hsp83* presumed promoter and P10 terminator driving expression of the DsRed fluorescent marker. The construct also contains the OpIE2:eGFP transformation marker and piggyBac arms for random integration. **B)** Fluorescence and brightfield images of transgenic L3 larvae (top) and male adult (bottom) showing strong and ubiquitous expression of DsRed in transgenic mosquitoes.

### Supplementary Figure S2

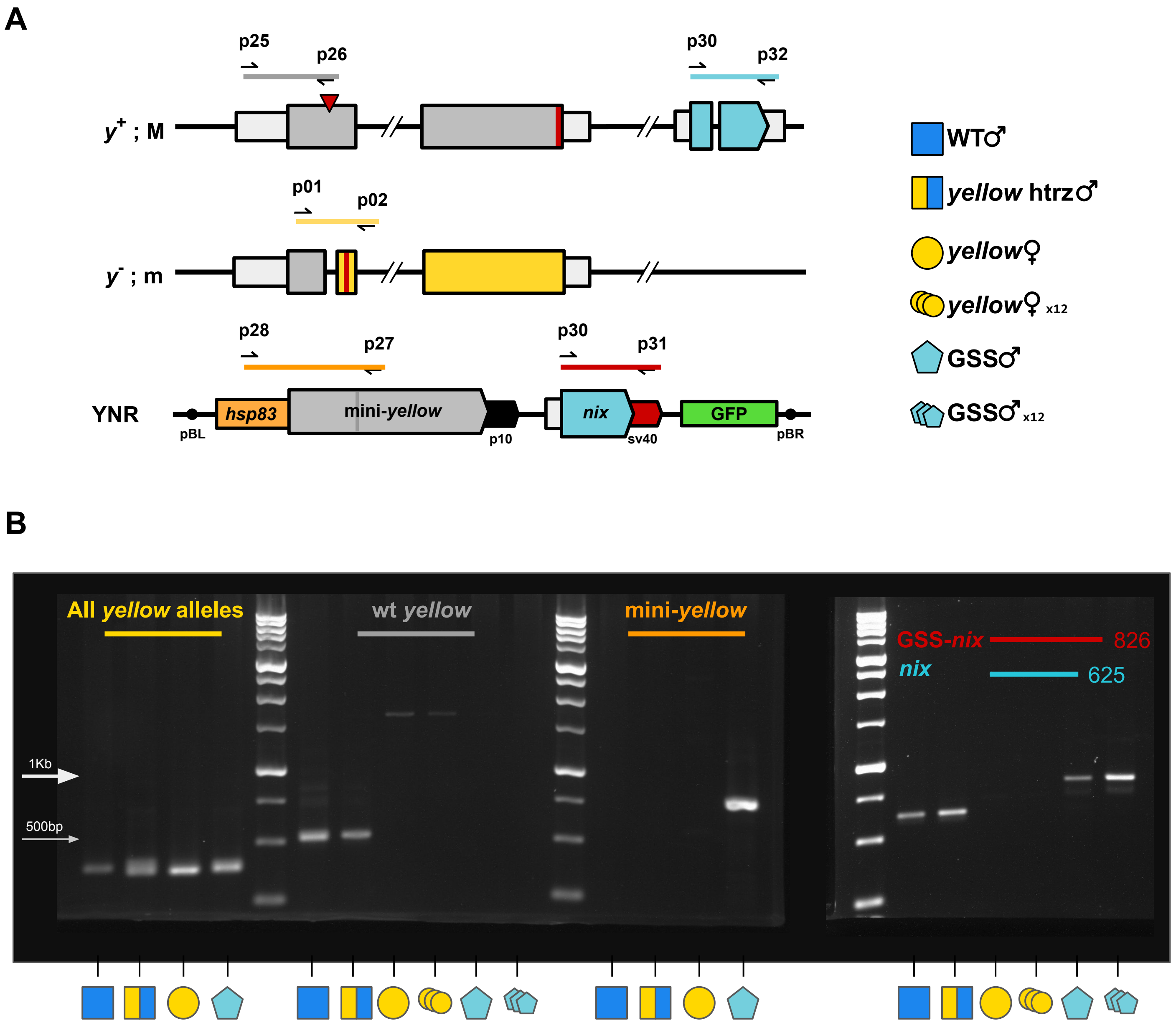


**Supplementary Figure 2. PCR genotyping of *yellow* and *Nix* alleles. A)** Schematic of three *yellow* alleles and the YRN construct with corresponding and discriminative PCR primers. A WT male chromosome ($y^+$; M) containing the native *yellow* gene and *nix* (~81 Mb upstream). A mutant female chromosome ($y^-$; m) lacking the *M* locus and containing a mutant *yellow* allele. The masculinizing mini-*yellow* YNR construct contains intron-less mini-*yellow* driven by the *hsp83* promoter and contains the *nix* 3&4 isoform from Lutrat et al 2022. Alleles or constructs can be differential detected - only WT *yellow* (grey), M-linked *nix* (teal), all *yellow* alleles (yellow), mini-*yellow* (orange), and GSS-*nix* (red). **B)** Gel electrophoresis of PCR products from single and pooled genomic DNA samples. Lanes are separated by a 1 kb ladder (GeneDirex, Taiwan).

### Supplementary Figure S3

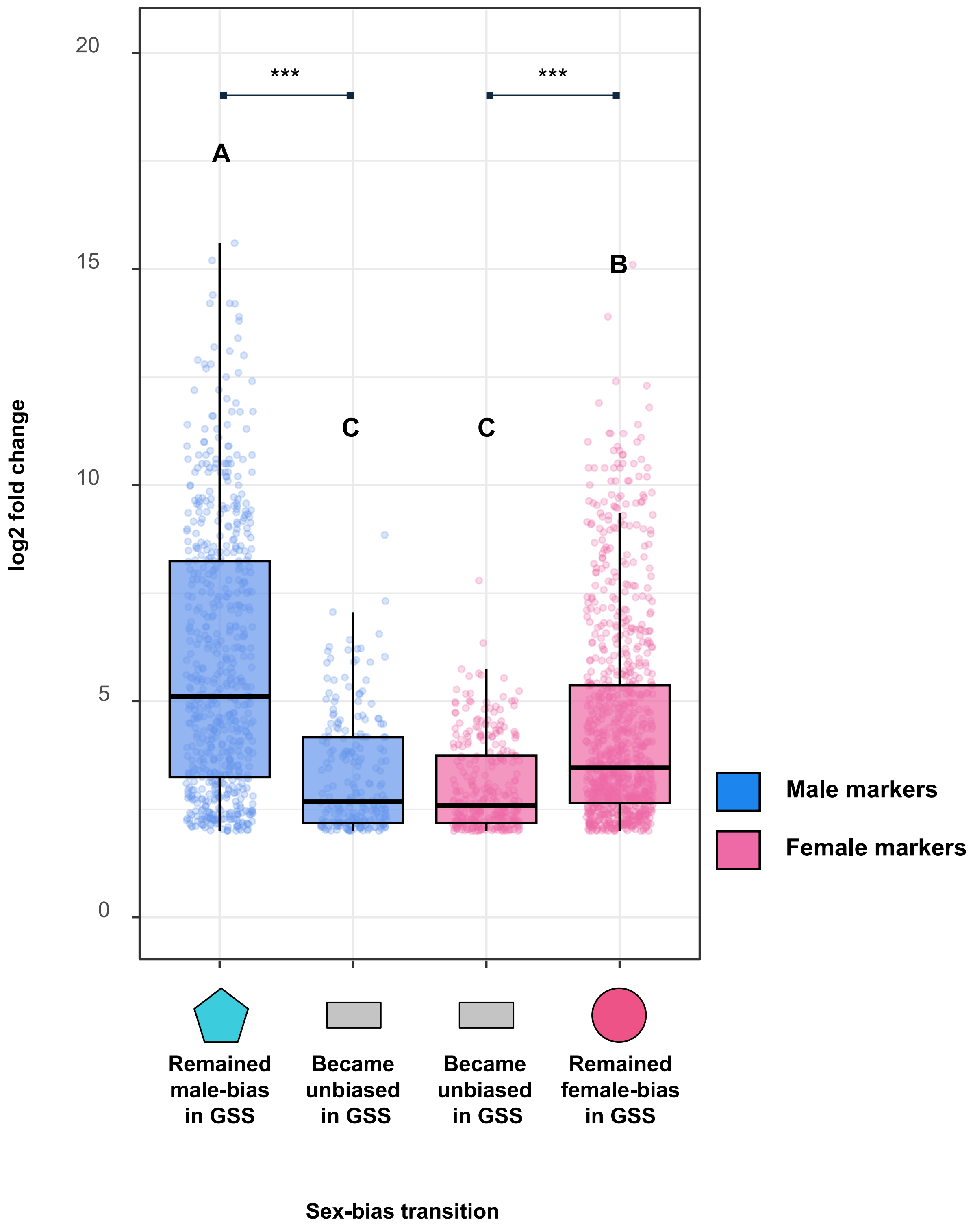


**Supplementary Figure 3. Magnitude of sex-biased expression in WTs for markers maintaining or losing sex-biased expression.** Log2 fold-change between the WT sexes for male-markers (blue) or females-markers (pink) that either maintained their sex-bias when comparing GSS pseudomales to WT females or became unbiased. Statistical significance within each sex was determined using a student's t-test (*** $p < 0.001$), and multiple comparisons across groups were performed using ANOVA followed by Tukey's HSD test. Significant differences are indicated by letter annotations ($p < 0.05$).

### Supplementary Figure S4

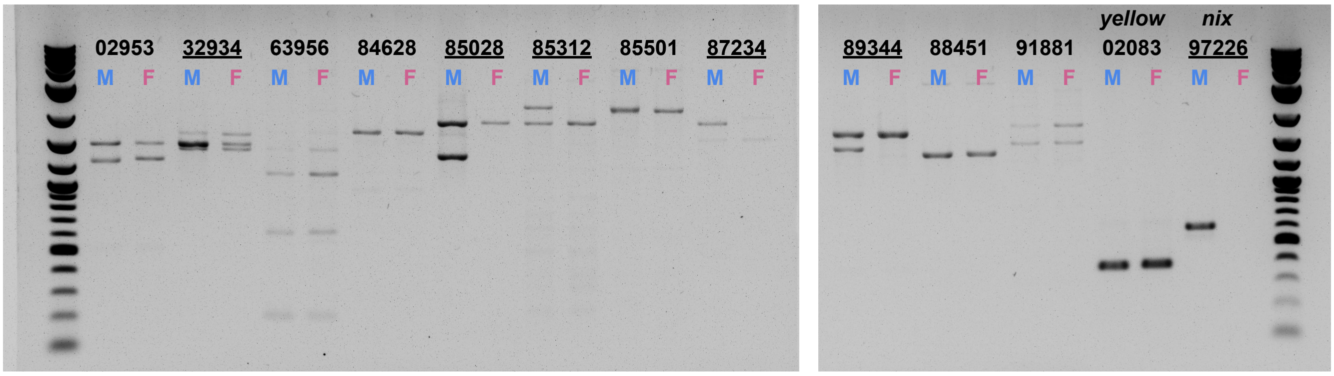


**Supplementary Figure 4. PCR-Based Validation of Male-Specific M-Locus Localization.** Gel electrophoresis of PCR amplicons from WT male (M) and female (F) genomic DNA, amplifying eleven male-biased genes whose expression is undetectable in GSS pseudomales. Amplification of *yellow* and *nix* are used as autosomal and M-specific controls, respectively. Gene IDs are indicated with the 5-digit LOC suffix; the left well shows the product from male DNA, and the right from female DNA. Underscored IDs indicate a male-specific amplification pattern.

### Supplementary Figure S5

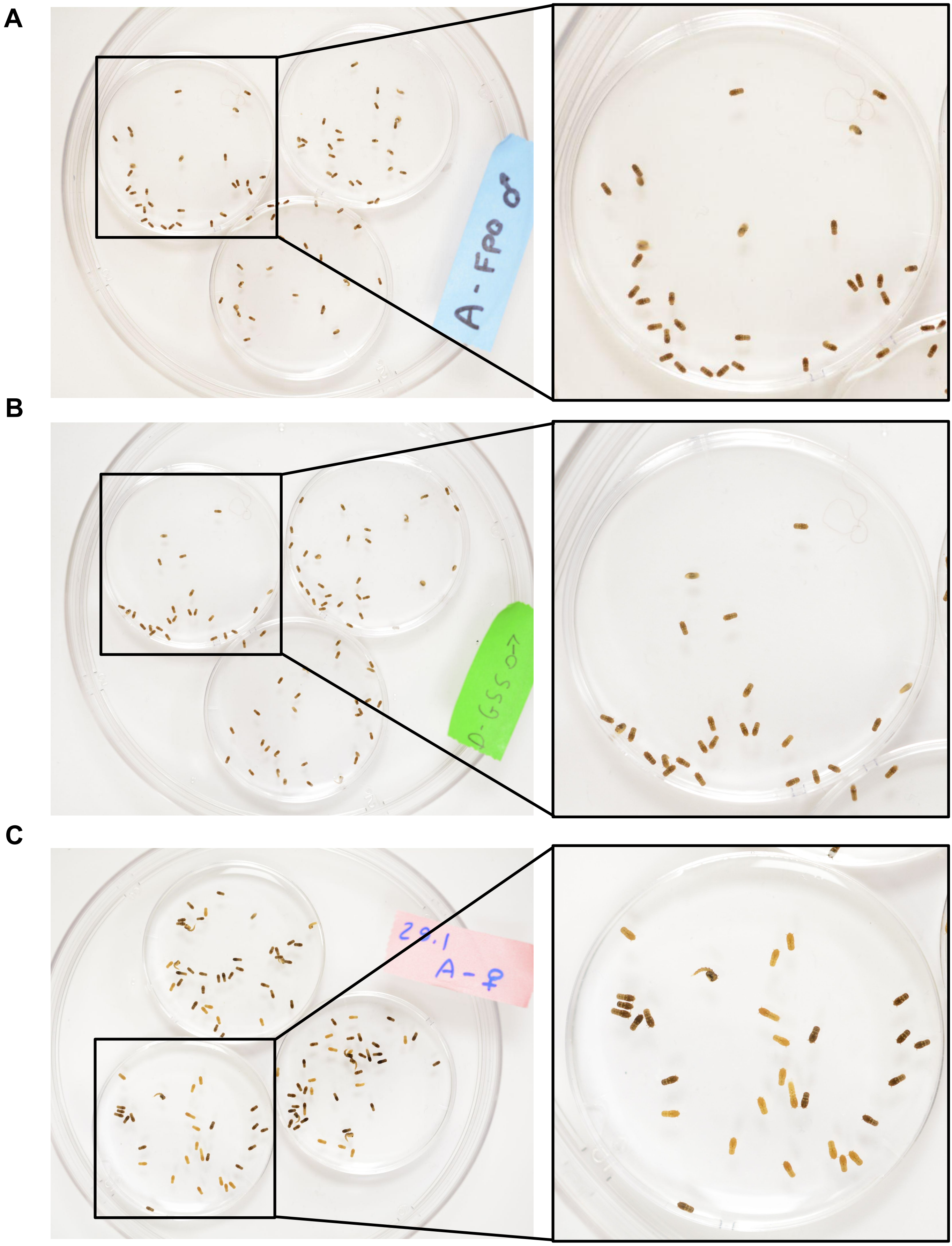


**Supplementary Figure 5. Imaging of pupal color and size.** Pupae from WT and GSS strains that were co-reared as larvae were sorted and photographed in three groups. **A)** WT males, **B)** GSS pseudomales, and C) females from both strains. Right panels show a high magnification photo of selected regions of the left image.

### Supplementary Figure S6

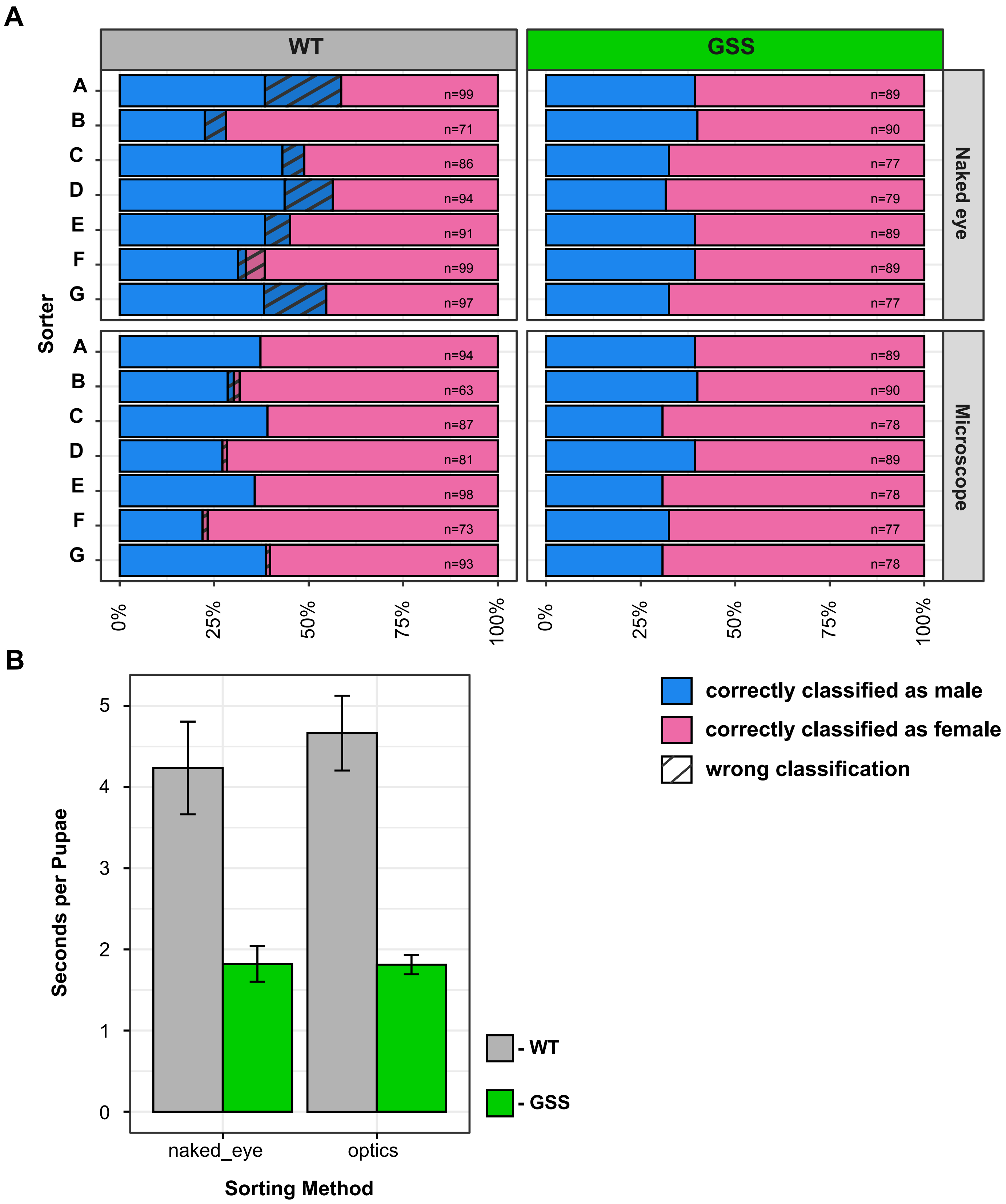


**Supplementary Figure 6. Benchmarking sex-separation for the GSS and WT strains A)** Seven researchers experienced in mosquito pupae sex separation sorters (A–G) each sorted pupae from the WT strain and *yellow*-GSS. For each strain, pupae were sexed either by the naked eye or with a dissecting microscope. The stacked bars show the proportion of pupae correctly classified as males (blue) or females (pink), with diagonal hatching indicating misclassifications. **B)** Speed of sex sorting for either method for the two strains (WT in gray and GSS in green). Error bars represent the standard error.
